# RNA polymerase II clustering through CTD phase separation

**DOI:** 10.1101/316372

**Authors:** M. Boehning, C. Dugast-Darzacq, M. Rankovic, A. S. Hansen, T. Yu, H. Marie-Nelly, D.T. McSwiggen, G. Kokic, G. M. Dailey, P. Cramer, X. Darzacq, M. Zweckstetter

**Author notes:** equal contribution, listed alphabetically (M. Boehning, C. Dugast-Darzacq, M. Rankovic). Correspondence should be addressed to (listed alphabetically): P.C.; X.D.; M.Z.

## Abstract

The carboxy-terminal domain (CTD) of RNA polymerase (Pol) II is an intrinsically disordered low-complexity region that is critical for pre-mRNA transcription and processing. The CTD consists of hepta-amino acid repeats varying in number from 52 in humans to 26 in yeast. Here we report that human and yeast CTDs undergo cooperative liquid phase separation at increasing protein concentration, with the shorter yeast CTD forming less stable droplets. In human cells, truncation of the CTD to the length of the yeast CTD decreases Pol II clustering and chromatin association whereas CTD extension has the opposite effect. CTD droplets can incorporate intact Pol II and are dissolved by CTD phosphorylation with the transcription initiation factor IIH kinase CDK7. Together with published data, our results suggest that Pol II forms clusters/hubs at active genes through interactions between CTDs and with activators, and that CTD phosphorylation liberates Pol II enzymes from hubs for promoter escape and transcription elongation.

---

Cellular processes often require clustering of molecules to facilitate their interactions and reactions^1,2^. During transcription of protein-coding genes, RNA polymerase (Pol) II clusters in localized nuclear hubs^3^. Whereas Pol II concentration in the nucleus is estimated to be ~1 μM, it increases locally by several orders of magnitude^4^. Such high Pol II concentrations are reminiscent of the clustering of proteins in membrane-less compartments such as P granules, Cajal bodies and nuclear speckles^1,2,5,6^. These cellular compartments are stabilized by interactions between intrinsically disordered low-complexity domains (LCD) and depend on liquid-liquid phase separation (LLPS) ^1,2,6–11^. However, the molecular basis of Pol II clustering remains unknown.

The largest subunit of Pol II, RPB1, contains a C-terminal low-complexity domain (CTD) that is critical for pre-mRNA synthesis and co-transcriptional processing^12^. The CTD is conserved from humans to fungi, but differs in the number of its hepta-peptide repeats with the consensus sequence Y_1_S_2_P_3_T_4_S_5_P_6_S_7_^13,14^. The human CTD (hCTD) contains a N-terminal half, which comprises 26 repeats and resembles the CTD from the yeast *Saccharomyces cerevisiae* (yCTD), and a C-terminal half containing 26 repeats of more divergent sequence (**Supplementary Fig. 1a**). CTD sequences from different species all contain a high number of tyrosine, proline and serine residues (**Supplementary Fig. 1b**)^13,15^. The most conserved CTD residues are Y_1_ and P_6_ that are present in all 52 repeats of hCTD. Truncation of the CTD of RPB1 in *Saccharomyces cerevisiae* to less than 13 repeats leads to growth defects and a minimum of eight repeats is required for yeast viability^16^. The CTD forms a mobile, tail-like extension from the core of Pol II^14^ that is thought to facilitate the binding of factors for co-transcriptional RNA processing and histone modification^13,14^.

Despite its extremely high conservation, its essential functions, and a large number of related published studies, the unique CTD structure and properties have remained enigmatic. Here we show that the CTD can undergo cooperative liquid-liquid phase separation *in vitro* that is driven by weak multivalent interactions. We further show that the CTD is critical for the formation of hubs of Pol II in human cells. Together with published results, we arrive at a model for gene activation that involves CTD-mediated Pol II clustering at active gene promoters, and release of initiated polymerases from these clusters after CTD phosphorylation.

## RESULTS

### CTD of Pol II phase separates into liquid-like droplets

To investigate whether liquid-liquid phase separation of CTD may underlie Pol II clustering, we expressed and purified hCTD and yCTD from *Escherichia coli*. We used a prokaryotic expression system to prevent eukaryotic post-translational modifications. The biophysical properties of the purified CTD proteins were characterized using circular dichroism (CD; **Supplementary Fig. 1c**). CD spectroscopy showed that hCTD and yCTD are intrinsically disordered in solution (**Supplementary Fig. 1c**) ^17–20^, consistent with the low complexity of CTD sequences (**Fig. 1a**) ^14^.

**Figure 1.**
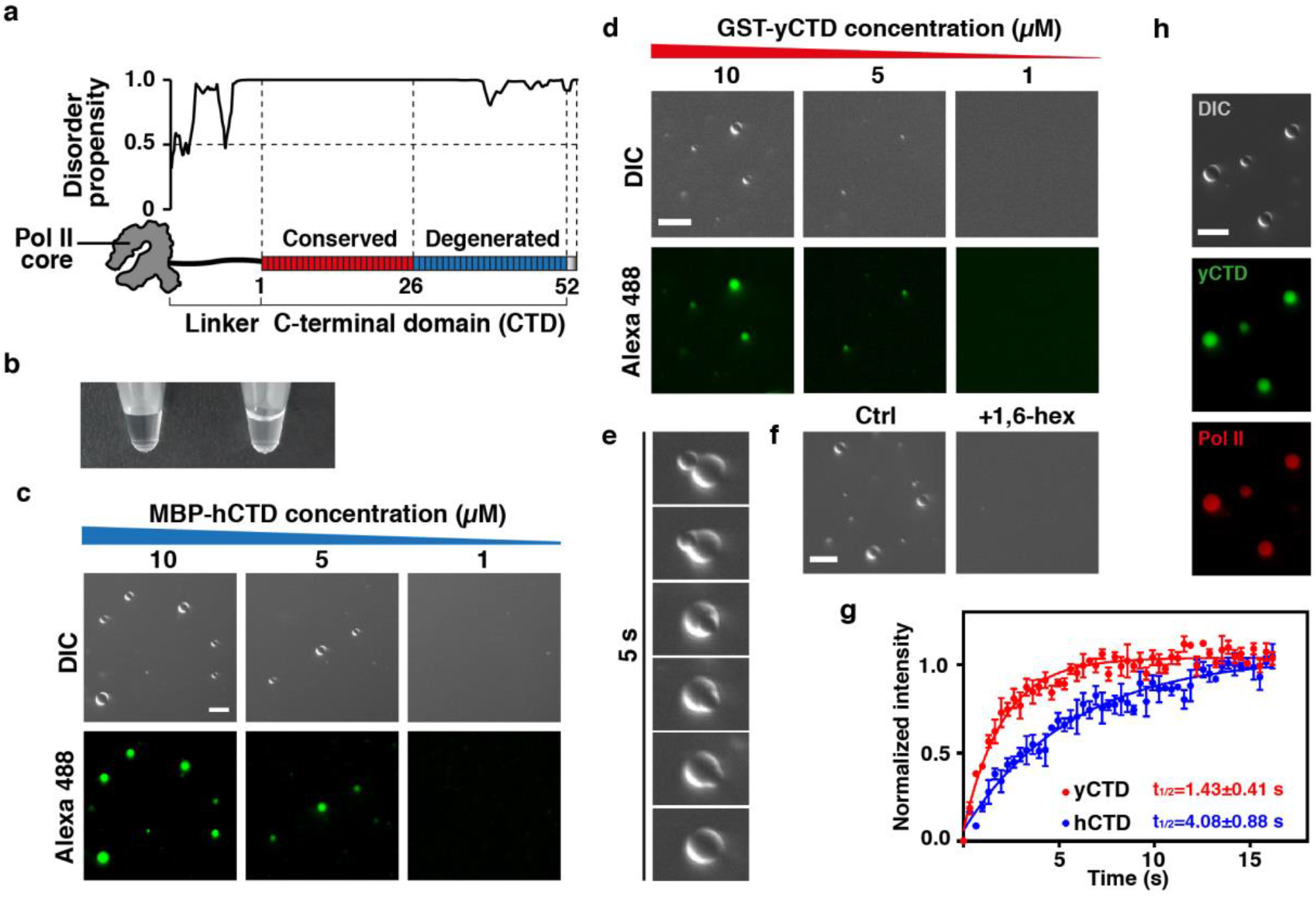
Phase separation of Pol II CTD into liquid-like droplets. (**a**) Disorder analysis (top) and schematic view of the low-complexity human C-terminal domain (hCTD) sequence of Pol II, comprising 52 conserved hepta-repeats (bottom). Its N-terminal half (red) is composed almost exclusively of consensus repeats (Y_1_S_2_P_3_T_4_S_5_P_6_S_7_) and is highly similar to the CTD in yeast (yCTD). (**b**) Addition of 16% of the molecular crowding agent dextran to a 20 μM solution of MBP-hCTD turns the solution turbid, a characteristic property of liquid phase separation. (**c**) DIC and fluorescence microscopy demonstrate the concentration-dependent formation of liquid droplets of MBP-hCTD in the presence of 16% dextran. (**d**) Concentration-dependent liquid phase separation of GST-yCTD in the presence of 16% dextran. (**e**) GST-yCTD droplets rapidly fuse upon contact into one spherical droplet (25 μM GST-yCTD in 16% dextran). (**f**) Liquid phase separation of yCTD is sensitive to the aliphatic alcohol 1,6-hexanediol (10 %). (**g**) FRAP kinetics of photobleaching a spot within hCTD (blue) and yCTD (red) droplets, which were formed in the presence of 16% dextran. The MBP fusion-tag was cleaved with TEV protease from MBP-hCTD and MBP-yCTD prior to the FRAP measurements. Data points represent mean values across three independent replicates and error bars show the standard error. (**h**) Pol II (red, Alexa 594) is concentrated in preformed yCTD droplets (green, Alexa 488). Scale bars are 10 μm in all panels.

Next we investigated the ability of CTD to undergo liquid-liquid phase separation using a combination of differential interference contrast (DIC) microscopy and fluorescence microscopy. DIC microscopy revealed the formation of micrometer-sized droplets at a concentration of 20 μM hCTD in the presence of 5-10% of the molecular crowding agent dextran (**Supplementary Fig. 2a**). Fluorescence microscopy demonstrated that hCTD molecules were strongly concentrated within the droplet interior compared to the surrounding milieu (**Supplementary Fig. 2a, lower panels**). At higher dextran concentration (16%), droplets could be detected already at a concentration of 5 μM hCTD (**Fig. 1b-c**). The number of droplets increased with increasing protein concentration (**Fig. 1c**), consistent with the general concentration-dependence of liquid phase separation^21^. In addition, hCTD formed droplets in the presence of another molecular crowding agent, the polysaccharide Ficoll (**Supplementary Fig. 2b**). hCTD also underwent LLPS after cleavage of the maltose-binding protein (MBP)-tag, while MBP alone did not form droplets in presence of molecular crowding agents (**Supplementary Fig. 2c**). hCTD droplet formation was robust against changes in ionic strength (**Supplementary Fig. 2d**), and incubation of the sample for one hour at different temperatures (**Supplementary Fig. 2e**). Like hCTD, yCTD formed droplets in a concentration-dependent manner (**Fig. 1d; Supplementary Fig. 3a-b**). Contacts of both hCTD and yCTD droplets led to fusion and formation of a single spherical droplet (**Fig. 1e; Supplementary movie 1 and 2**). At concentrations subcritical for LLPS, yCTD was incorporated into preformed hCTD droplets and hCTD was included into preformed yCTD droplets (**Supplementary Fig. 3c**), in agreement with the ability of the CTD to be trapped into droplets and hydrogels of LCD proteins ^22,23^. Formation of yCTD droplets was also resistant against changes in ionic strength (**Supplementary Fig. 3d**) and temperature (**Supplementary Fig. 3e**), similar to hCTD. The combined data show that the CTD of Pol II forms LCD-LCD interactions and readily undergoes LLPS to form liquid-like droplets in solution.

Liquid droplets and cellular puncta are held together by weak, distributed interactions between LCDs that are sensitive to aliphatic alcohols^6,24,25^. As expected for such interactions, liquid phase separation of yCTD and hCTD was counteracted by addition of 5-10% 1,6-hexanediol (**Fig. 1f; Supplementary Fig. 4a-b, upper panels**). Addition of 5-10% of the hexanediol isomer 2,5-hexanediol also inhibited CTD droplet formation (**Supplementary Fig. 4a-b, lower panels**). Because it was shown that 2,5-hexanediol is less efficient in dissolving droplets and hydrogels^26^, the data indicate that CTD droplets are more sensitive to aliphatic alcohols than other LCD-LCD interactions. On the contrary, CTD phase separation is robust to changes in ionic strength (**Supplementary Fig. 2d and 3d**).

### CTD length influences CTD phase separation *in vitro*

A characteristic property of liquid-like droplets is fast diffusion of molecules in their interior^1^. We used fluorescence recovery after photobleaching (FRAP) to compare diffusion kinetics of hCTD and yCTD molecules within droplets. MBP-tagged hCTD and yCTD proteins were fluorescently labeled on a single cysteine residue that is present C-terminal to the TEV protease cleavage site. After cleavage of the MBP tag and droplet formation, circular regions in the interior of CTD droplets were bleached. Within hCTD droplets, the bleached fluorescence recovered with a half time of 4.08 s ± 0.88 s (**Fig. 1g**). For yCTD, recovery was faster with a half time of 1.43 s ± 0.41 s. (**Fig. 1g**).

These results demonstrate that CTD molecules within droplets are generally highly dynamic, confirming the liquid-like nature of CTD droplets. The difference in fluorescence recovery between hCTD and yCTD further suggests that the higher number of repeats in hCTD strengthens CTD-CTD interactions. This observation is consistent with the concentration-dependent ability of hCTD and yCTD to undergo LLPS when fused to the MBP-tag. MBP-hCTD phase separated at a concentration of 5 μM (**Fig. 1c; Supplementary Fig. 5, top**). In contrast, LLPS of MBP-yCTD started only at a 4-6 fold higher protein concentration (**Supplementary Fig. 5, middle**). When the smaller, dimerizing glutathione S-transferase (GST)-tag was used to replace the more soluble MBP-tag^27^, the critical concentration for yCTD phase separation decreased to approximately 5 μM (**Supplementary Fig. 5, bottom;** **Fig. 1d**). These results suggest that the solubilizing effect of MBP counteracts droplet formation. This effect is more easily overcome by hCTD because the higher repeat number and valency results in stronger CTD-CTD interactions compared to yCTD. We conclude that the length of the CTD influences the stability and dynamics of LLPS droplets, with a longer CTD leading to stronger CTD-CTD interactions and less dynamic droplets.

### CTD droplets recruit intact Pol II

The above results indicate that CTD-CTD interactions within liquid droplets may underlie Pol II clustering. However, we could not test directly whether intact Pol II forms LLPS droplets *in vitro* because it was impossible to prepare Pol II at a sufficient concentration in the presence of dextran or Ficoll. We could however test whether Pol II could be trapped within CTD droplets. We purified Pol II from yeast cells, labeled it with the fluorescent dye Alexa 594 and added it to pre-formed CTD droplets at a concentration of 0.02 μM. Fluorescence microscopy showed that Pol II located to CTD droplets (**Fig. 1h**).

### CTD length controls Pol II clustering in human cells

To explore whether CTD-based LLPS may underlie Pol II clustering in cells, we engineered two human cell lines that express a fluorescent Dendra2-tagged version of RPB1. To create these cell lines we transfected cells with a plasmid containing an α-amanitin resistant RPB1 variant (N792D) and selected cells in the presence of α-amanitin, which leads to the degradation of endogenous RPB1^3^. Such cell lines are known to recapitulate the behavior of endogenous wild-type Pol II^3,28,29,30^. One cell line contained the full-length CTD with 52 repeats (RPB1-52R), whereas the other cell line contained a truncated CTD with 25 repeats (RPB1-25R) that closely resembles the yCTD sequence (**Fig. 2a**). The two cell lines remained viable upon degradation of endogenous RPB1 after treatment with α-amanitin and expressed similar levels of the Dendra2-tagged exogenous Pol II as assessed by western blotting (**Fig. 2b**), confocal imaging and fluorescence-activated cell sorting (FACS; **Supplementary Fig. 6a-c**). The two cell lines also grew at a similar rate (**Supplementary Fig. 6d**).

**Figure 2.**
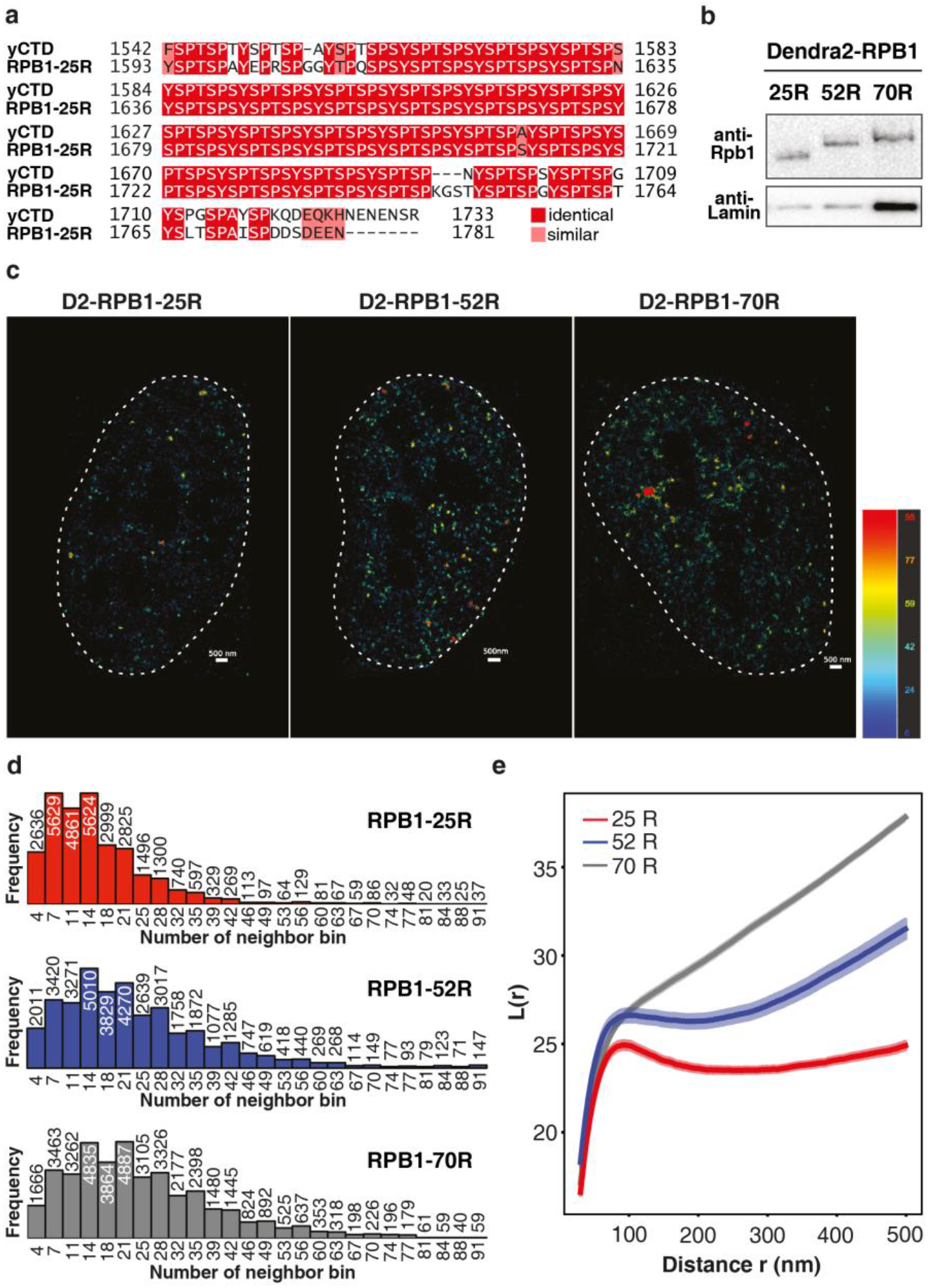
CTD-dependent Pol II clustering in human cells. (**a**) Pairwise alignment of CTD sequences from the yeast *S. cerevisiae* and the human Dendra2-RPB1-25R cell line. The RPB1-25R cell line encodes a truncated version of human RPB1 comprising only 25 CTD repeats (Methods). This hCTD truncation closely resembles the yeast CTD in length (189 aa (RPB1-25R) vs. 192 aa (yCTD)) and amino acid composition (similarity (identity): 87.2% (83.2%)). The alignment was generated using the EMBOSS needle tool^43^ with default settings and aligned residues are colored in different shades of red according to their similarity (consensus symbol). (**b**) Western blot analysis of the expression level of Dendra2-RPB1-25R, 52R and 70R. The RPB1-70R is expressed at lower level than the other two proteins. (**c**) 3D-PALM reconstruction of RPB1-25R (left), RPB1-52R (middle) and RPB1-70R (right). Each detection is color-coded by the number of detections lying in a radius of 120 nm around it. The color scale is: number of detection / disk (120 nm). (**d**) Local density distribution (radius = 120 nm). Histograms of the average number of detections in a 120 nm-radius disk of Dendra2-RPB1-25R (top), Dendra2-RPB1-52R (middle) and Dendra2-RPB1-70R (bottom). (**e**) *L*-modified Ripley function. Linearized representation of the classic Ripley function. The null model of complete spatial randomness is rejected since the curves positively deviate from zero. All three curves exhibit strong clustering at all scales.

We now studied clustering of Pol II in these human cell lines with the help of 3D-PALM super-resolution microscopy using induced astigmatism by a cylindrical lens (**Fig. 2c-e; Supplementary Fig. 6e**)^3,31^. Compared to cells with full-length CTD (52R), cells with the truncated, yeast-like CTD (25R) showed less Pol II clustering (**Fig. 2c-d**). These results suggested that CTD interactions underlie Pol II clustering in cells and that the CTD length influences clustering. To test this directly, we further created a cell line containing an artificially extended CTD (RPB1-70R, Online Methods). This cell line was also viable and grew at a similar rate as the other two lines upon degradation of endogenous RPB1 (**Supplementary Fig. 6d**), though it expressed RPB1 at a lower level (**Fig. 2b**). Despite this difference in expression level, the 70R cell line showed even more Pol II clustering than cells with wild-type, full-length CTD (**Fig. 2c-d**), strongly supporting our findings.

For all three cell lines, differences in CTD-dependent cluster density were supported by quantitative analysis on the basis of a modified Ripley function *L(r)*, which compares the spatial distribution of localizations to complete spatial randomness *(L(r)*=0 for all r) ^32^. In all cells, *L(r)* curves showed a strong clustering signature (**Fig. 2e; Supplementary Fig. 6e**). Whereas the sharp increase observed at scales less than 100 nm can be influenced by photophysical effects, such as blinking of Dendra2^33^, the continuous increase at larger spatial distances is representative of Pol II clustering at multiple length scales. Taken together, these results demonstrate that Pol II clustering in cells depends on the CTD and increases with increasing CTD length.

### CTD length influences Pol II dynamics in cells

We next investigated the impact of CTD length on Pol II dynamics *in vivo* using two orthogonal approaches, live-cell single particle tracking (SPT)^34^ and FRAP experiments. Because these methods require a high signal-to-noise ratio and a photo-stable fluorescent label, we established cell lines with a Halo-tag on RPB1 containing 25, 52 and 70 CTD repeats (25R, 52R and 70R, respectively) (**Supplementary Fig. 7**). We then tracked single molecules of Pol II in live cells as demonstrated by single-step photo-activation and -bleaching (**Fig. 3a-b**, **Supplementary movies 3-5**). Subsequent two-state kinetic modeling analysis assuming a free and bound state (**Fig. 3c**, **Supplementary Fig. 8a**) revealed that 29.1% of wild-type Pol II (RPB1-52R) in live cells was immobile and therefore presumably chromatin-associated. The bound Pol II fraction was decreased to 21% in RPB1-25R cells and was increased to 38.4 % in RPB1-70R cells (**Fig. 3d; Supplementary Fig. 8b-c**). In addition, the diffusion coefficients for free Pol II were higher and lower, respectively, for RPB1-25R and RPB1-70R cells. Free diffusion coefficients of 3.74, 2.97, and 2.34 μm^2^/s were measured in RPB1-25R, RPB1-52R and RPB1-70R cells, respectively (**Fig. 3e**). These large differences in diffusion coefficients cannot be explained solely by differences in mass or size (Online Methods). Therefore, our results indicate that CTD length strongly influences Pol II mobility *in vivo*, with shorter and longer CTDs leading to higher and lower mobility, respectively.

**Figure 3.**
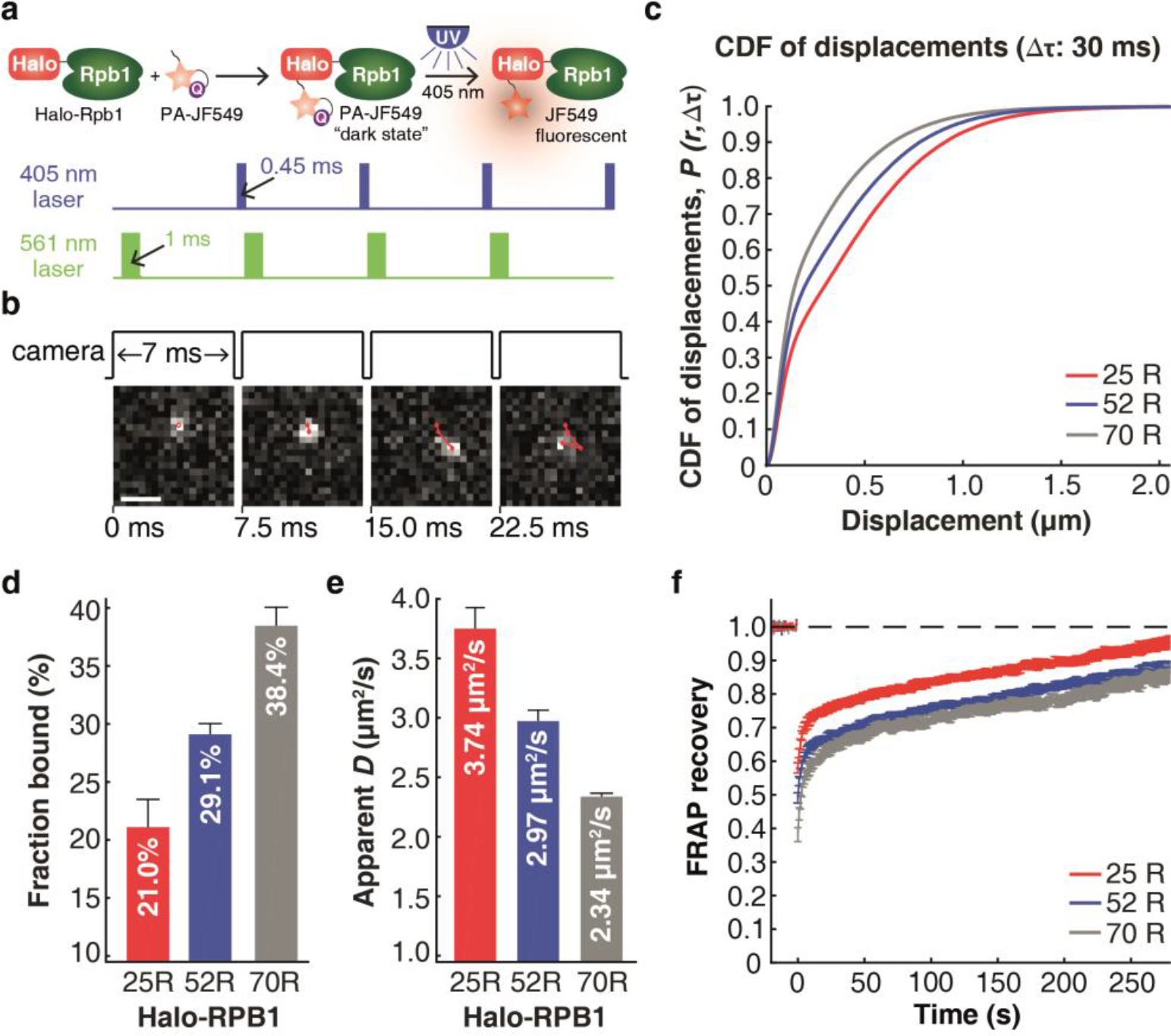
CTD-dependent Pol II dynamics in human cells (**a**) Overview of spaSPT at ~133 Hz. Halo-RPB1 labeled with PA-JF_549_ is photo-activated with a 405 nm laser and excited with 1 ms stroboscopic pulses of a 561 nm laser. This simultaneously minimizes motion-blurring by strobing the excitation laser and minimizes tracking errors by maintaining a low average density of ~1 localization per frame. (**b**) Representative spaSPT images with overlaid trajectories. The scale bar corresponds to 1 μm. (**c**) Cumulative distribution functions (CDF) for displacements. The CDF of displacements for the representative time-lag Δτ = 30 ms is shown for Halo-RPB1-25R, Halo-RPB1-52R and Halo-RPB1-70R. The data shown is merged from three independent replicates. (**d**) Bound fractions of 25R-, 52R- and 70R-Halo-RPB1. The bound fraction was inferred from two-state model fitting to the spaSPT displacement data using Spot-On^34^. Each of three independent replicates was fitted separately and bar graphs show the mean and standard error. (**e**) Diffusion coefficients of the free population of 25R-, 52R- and 70R-Halo-RPB1. Free diffusion coefficients were inferred from two-state model fitting to the spaSPT displacement data using Spot-On^34^. Each of three independent replicates was fitted separately and bar graphs show the mean and standard error. (**f**) FRAP dynamics. Mean drift and photobleaching-corrected FRAP recoveries are shown for Halo-RPB1-25R, Halo-RPB1-52R and Halo-RPB1-70R. FRAP data were collected at 1 frame per second for 300 seconds and bleaching was performed before frame 21. FRAP curves show mean across three independent replicates and error bars show the standard error.

These findings in cells match our observed length-dependence of CTD-CTD interactions *in vitro* (**Fig. 1g; Supplementary Fig. 5**). Indeed, FRAP recovery curves in human cells depended on CTD length (**Fig. 3f**), consistent with differences in FRAP recovery kinetics observed between hCTD and yCTD droplets *in vitro* (**Fig. 1g**). Analysis of these FRAP recovery curves by a reaction-dominant two-state model^35,36^ further showed that the fraction that did not recover within a few seconds increased from 27% in RPB1-25R cells to 35% in RPB1-52R cells to 38% in RPB1-70R cells (**Supplementary Fig. 8d-f**). This trend is consistent with the SPT results (**Fig. 3d**), which also showed a higher chromatin-associated fraction for Pol II with a longer CTD. Interestingly, both SPT and FRAP analysis showed that this putative chromatin-associated fraction of RNA Pol II was decreased to similar levels in all three cell lines after flavopiridol treatment, which blocks the transition into productive elongation by targeting P-TEFb (**Supplementary Fig. 9**). This favors an interpretation in which the CTD length dependent bound fraction is linked to polymerase activity. Together our data show that longer CTDs result in more clustered Pol II and chromatin association *in vivo*, reflecting the influence of CTD length on liquid-liquid phase separation *in vitro*.

### CTD phosphorylation dissolves droplets

Finally, we investigated whether CTD phosphorylation impacts phase separation. It has long been known that assembly of the pre-initiation complex at Pol II promoters requires an unphosphorylated CTD, and that subsequent CTD phosphorylation at S_5_ CTD residues by the cyclin-dependent kinase 7 (CDK7) in transcription factor IIH (TFIIH) stimulates the transition of Pol II into active elongation^37,38^. We treated hCTD with recombinant human TFIIH subcomplex containing CDK7 kinase^39^ and adenosine triphosphate (ATP), leading to S_5_ phosphorylation of hCTD (**Fig. 4a; Supplementary Fig. 10a**). The resulting CDK7-phosphorylated hCTD was no longer able to form droplets, whereas prior incubation with ATP alone did not inhibit LLPS (**Fig. 4b**). Phosphorylation of yCTD by the yeast TFIIH kinase subcomplex also inhibited phase separation (**Supplementary Fig. 10b-c**). In addition, phosphorylation of preformed hCTD droplets by human CDK7 caused gradual shrinking and ultimately disappearance of hCTD droplets (**Fig. 4c; Supplementary movie 6**). Therefore, phosphorylation at S5 positions is incompatible with CTD phase separation and transfers the CTD from the highly concentrated state within droplets to the dispersed pool.

**Figure 4.**
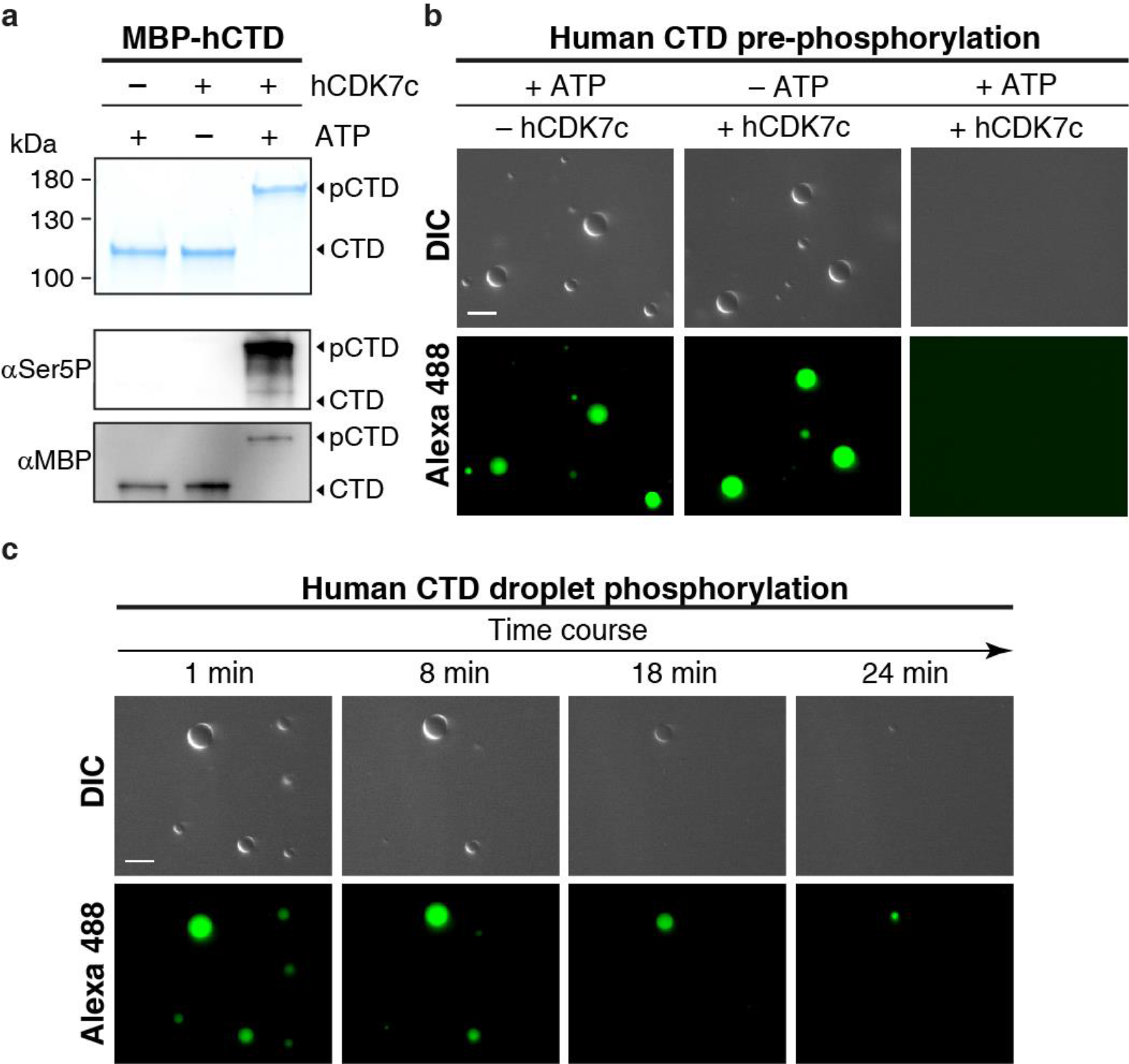
Liberation through initiation-coupled CTD phosphorylation. (**a**) SDS-PAGE and western blot analysis of phosphorylated MBP-hCTD fusion protein. MBP-hCTD was treated with recombinant human CDK7 complex. The hCTD substrate became highly phosphorylated, resulting in a pronounced mobility change during polyacrylamide electrophoresis in comparison to the non-phosphorylated substrate (-ATP and ‒kinase control reactions). Western blotting confirms phosphorylation of heptad position Ser5. Corresponding loading controls are shown to correct for potential differences in blotting efficiency. (**b**) hCTD phase separation is inhibited upon CTD phosphorylation by the human TFIIH subcomplex containing the CDK7 kinase. This effect is neither caused by hydrotropic properties of ATP^44^ nor the pure presence of the kinase, since MBP-hCTD readily forms droplets in control reactions containing ATP or the kinase alone. (**c**) CDK7 phosphorylation dissolves preformed hCTD droplets. Scale bars correspond to 10 μm.

## DISCUSSION

Here we show that the Pol II CTD can undergo length-dependent liquid-liquid phase separation *in vitro*, and that it controls Pol II clustering and mobility *in vivo*. Whereas CTD function is generally thought to depend on defined binary interactions of short CTD regions (1-3 repeats) with CTD-binding proteins, our results suggest that CTD function can additionally depend on weak homo- and heterotypic LCD-LCD interactions, and that these interactions may dominate Pol II localization and dynamics *in vivo*. Although it was previously described that the CTD can interact with pre-formed LLPS droplets and hydrogels of FET proteins ^22,23^, our results demonstrate for the first time that the CTD alone, in absence of other proteins, can undergo phase separation. Whereas the correlation between our *in vitro* and *in vivo* data is striking, our data cannot demonstrate directly that RNA polymerase II undergoes phase separation *in vivo*. However, we provide strong evidence that the weak and multivalent interactions driving phase separation *in vitro* are the same that drive polymerase hubs within the nucleoplasm of living cells.

Our findings have implications for understanding Pol II transcription in eukaryotic cells and suggest a simple model for gene activation and the initiation-elongation transition during early transcription (**Fig. 5**). Unphosphorylated Pol II clusters forming nucleoplasmic hubs in cells, mediated by CTD-CTD interactions. Pol II hubs may be recruited by transcriptional activators that bind to regulatory sites such as enhancers^40^. Transcriptional activators can also undergo LCD interactions^41^ and might assist in Pol II hub formation when Pol II concentration is subcritical. Formation of Pol II hubs near gene promoters may provide a reservoir of Pol II to achieve high initiation rates during activated transcription. When a Pol II enzyme is incorporated into a preinitiation complex, its CTD gets phosphorylated by the TFIIH kinase CDK7. Phosphorylation removes this Pol II enzyme from the hub, liberates it to escape the promoter and enables transition into active transcription elongation (**Fig. 5**).

**Figure 5.**
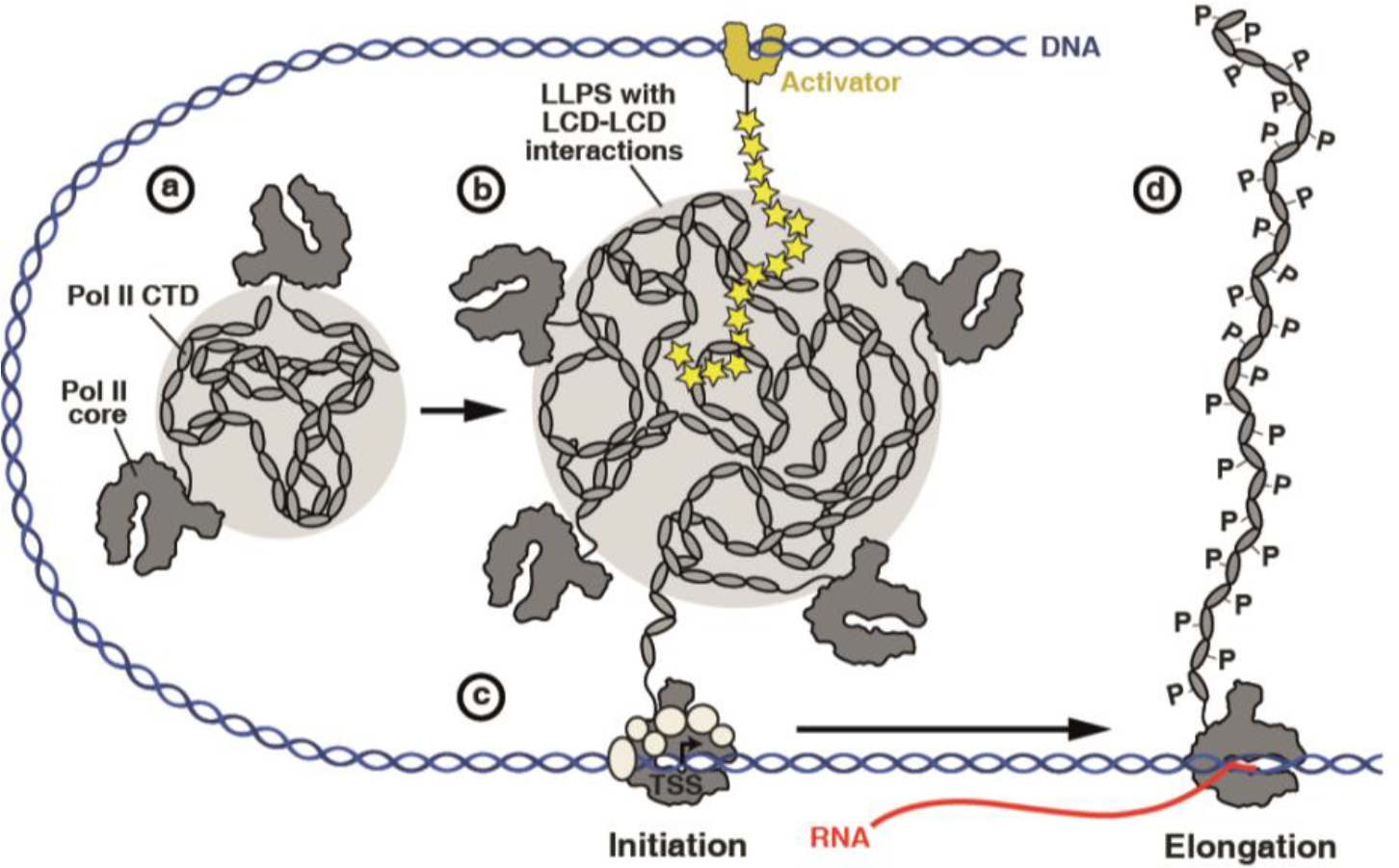
Model for the role of CTD-driven phase separation in activated transcription. Activators may recruit or nucleate Pol II hubs near gene promoters. Initiation-coupled CTD phosphorylation removes individual Pol II enzymes for transcription elongation.

When our manuscript was under review, the Zhou laboratory published that phase separation of the intrinsically disordered region of cyclin T1, which is a subunit of the positive elongation factor P-TEFb, promotes inclusion of RNA polymerase II CTD into hubs ^42^. This effect was observed exclusively for the CTD in its CDK7-phosphorylated, but not its unphosphorylated form^42^. Together with the current work, this suggests that the Pol II CTD can undergo phase separation via at least two different mechanisms: in its unphosphorylated form, CTD phase separation is based mainly on weak hydrophobic interactions (**Fig. 1f, Supplementary Fig. 4**). Phosphorylation disrupts CTD-CTD interactions (**Fig. 4**) and successively allows the CTD to engage in novel multivalent interactions with selected factors. The latter interactions are likely primarily electrostatic in nature and can also lead to phase separation, as demonstrated for the phosphorylated CTD together with the positively charged intrinsically disordered region of cyclin T1^42^. Our results and those of Lu et al.^42^ are thus complementary and provide a starting point for analyzing the chemical basis for CTD phase separation, its possible modulation by nucleic acids and protein factors, and its specific roles in transcription regulation and the coordination of the transcription cycle. More generally, this provides a powerful and highly specific and regulated local protein sorting mechanism modulating local proteomes within cells.

## METHODS

Methods and any associated references are available in the online version of the paper.

*Note: Any Supplementary Information and Source Data files are available in the online version of the paper*.

## ACKNOWLEDGMENTS

We thank Seychelle M. Vos for the MBP-hCTD expression plasmid and advice on purification, Sandra Schilbach for the yeast TFIIH kinase module as well as Andrea Boltendahl and Carina Burzinski for help with cloning and purification. We are grateful to Susmitha Ambadipudi for help with initial microscopy measurements and discussions about LLPS. We thank Luke Lavis for generously providing JF dyes. We thank Anatalia Robles, Carla Inouye, Shuang Zheng, Mallory Haggart and Janeen Lim for technical and administrative assistance. In vivo FRAP experiments were conducted at the CRL Molecular Imaging Center, supported by Gordon and Betty Moore Foundation. We thank all current members of the Cramer, Zweckstetter and Tjian-Darzacq labs for discussions and Robert Tjian for critical reading of the manuscript. M.Z. was supported by the Deutsche Forschungsgemeinschaft (SFB860; project B02), by the Cluster of Excellence and DFG Research Center Nanoscale Microscopy and Molecular Physiology of the Brain and the advanced grant ‘LLPS-NMR’ of the European Research Council. P.C. was supported by the Deutsche Forschungsgemeinschaft (SFB860; project A13), the advanced grant ‘TRANSREGULON’ of the European Research Council, and the Volkswagen Foundation. X.D. was supported by NIH grant UO1-EB021236 and the California Institute of Regenerative Medicine grant LA1-08013.

## AUTHOR CONTRIBUTIONS

M.B. designed experiments, generated constructs and prepared proteins unless otherwise noted. C. D-D designed experiments and established and characterized the RPB1 cell lines and performed and analyzed the *in vivo* FRAP and SPT experiments. M.R. designed experiments, performed all phase separation assays, DIC and fluorescence microscopy, *in vitro* FRAP measurements and data analysis. A.S.H. designed, performed and analyzed SPT experiments and helped with the *in vivo* FRAP analysis. H.M.N. designed, performed and analyzed 3D PALM experiments. D.T.McS. performed cell viability experiments and helped in performing 3D PALM experiments. G.D. designed and cloned the different RPB1 expression vectors. G.K. prepared human TFIIH kinase complex. T.Y. performed CD and NMR experiments. C.D.-D., X.D., P.C. and M.Z. designed and supervised research. M.B., M.R., C.D.-D., P.C., X.D. and M.Z. prepared the manuscript with input from all authors.

## ONLINE METHODS

**Cloning and protein expression.** A plasmid encoding the human Pol II CTD sequence (hCTD) (RPB1 residues 1593-1970) fused C-terminally to a sequence encoding 6xHis-tagged maltose-binding protein (MBP) directly followed by a flexible linker (10xAsn) and a tobacco etch virus (TEV) protease cleavage site was a kind gift from Seychelle M. Vos (Max Planck Institute for Biophysical Chemistry, Göttingen). Derivative constructs, in which the hCTD sequence was replaced by the sequence coding for the *S. cerevisiae* Pol II CTD (yCTD) (RPB1 residues 1542-1733) or entirely removed, were generated using Gibson Assembly (New England Biolabs) and through deletion mutagenesis, respectively. All proteins contain a single cysteine residue C-terminal of the TEV protease cleavage site that allows for site-specific labeling. MBP-tagged proteins were overexpressed in *E. coli* BL21 (DE3) RIL cells (Stratagene) cultured in LB medium containing 50 mg/l kanamycin and 34 mg/l chloramphenicol. After reaching an OD600 of ~0.8, 0.5 mM IPTG was added and proteins were expressed for 3-4 h at 37 °C. Cells were harvested by centrifugation and resuspended in lysis buffer LB300 (20 mM HEPES pH 7.4, 300 mM NaCl, 30 mM imidazole, 10 % glycerol, 1 mM DTT, 0.284 μg/ml leupeptin, 1.37 μg/ml pepstatin A, 0.17 mg/ml PMSF, 0.33 mg/ml benzamidine). The cell suspension was snap frozen and stored at −80 °C.

The sequence coding for the yCTD was additionally cloned into a pET24-derived plasmid, directly C-terminal to a GST-tag followed by the TEV protease cleavage site. An N-terminal 6xHis-tag was introduced by site-directed mutagenesis. From the obtained plasmid, a second expression vector encoding only 6xHis-GST-TEV was constructed through deletion mutagenesis. GST-tagged proteins were overexpressed in *E. coli* BL21 Rosetta 2(DE3)pLysS cells (Stratagene) grown in 2xYT medium containing 50 mg/l kanamycin and 34 mg/l chloramphenicol. After the culture reached an OD_600_ of 0.6-0.8, Isopropyl β-D-1-thiogalactopyranoside (IPTG) was added to a final concentration of 0.5 mM. GST-yCTD was overexpressed for 16 h at 18 °C, GST for 3 h at 18 °C. Cells were harvested by centrifugation, resuspended in lysis buffer LB150 (20 mM HEPES pH 7.4, 150 mM NaCl, 30 mM imidazole, 10 % glycerol, 1 mM DTT, 0.284 μg/ml leupeptin, 1.37 μg/ml pepstatin A, 0.17 mg/ml PMSF, 0.33 mg/ml benzamidine), flash frozen in liquid nitrogen and stored at −80 °C.

Sequences encoding the full-length subunits of the human TFIIH kinase module (CDK7, MAT1 and cyclin H) were separately transferred into MacroBac 438B vectors^45^ and combined into a single construct by ligation independent cloning. All subunits contained a N-terminal 6xHis-tag followed by a TEV protease cleavage site. Insect cell expression was performed as described^46^.

**Protein purification.** All purification steps were performed at 4 °C. Frozen *E. coli* cell suspension was thawed, lysed by sonication, cleared from insoluble material by centrifugation (27,000 g, 45 min, 4 °C) and filtered through 0.8 μm syringe filters.

For the purification of MBP-tagged proteins, cleared *E. coli* lysate was loaded onto a 5 ml HisTrap HP column (GE healthcare) equilibrated in LB300. The HisTrap column was washed extensively using high salt buffer HSB1000 (20 mM HEPES pH 7.4, 1 M NaCl, 30 mM imidazole, 10 % glycerol, 1 mM DTT, 0.284 μg/ml leupeptin, 1.37 μg/ml pepstatin A, 0.17 mg/ml PMSF, 0.33 mg/ml benzamidine) and equilibrated again in LB300. The column was then attached in-line to a LB300-equilibrated XK-16 column (GE healthcare), which was packed with amylose resin (New England Biolabs). Bound proteins were eluted directly onto the amylose column using nickel elution buffer 300 (20 mM HEPES pH 7.4, 300 mM NaCl, 500 mM imidazole, 10 % glycerol, 1 mM DTT, 0.284 μg/ml leupeptin, 1.37 μg/ml pepstatin A, 0.17 mg/ml PMSF, 0.33 mg/ml benzamidine). The HisTrap column was subsequently removed and the amylose column was washed again extensively with HSB1000 buffer. MBP-tagged proteins were eluted using amylose elution buffer (20 mM HEPES pH 7.4, 300 mM NaCl, 10 % glycerol, 1 mM DTT, 117 mM maltose, 0.284 μg/ml leupeptin, 1.37 μg/ml pepstatin A, 0.17 mg/ml PMSF, 0.33 mg/ml benzamidine) and concentrated with a 30 kDa MWCO Amicon Ultra filter unit (Merck). The concentrate was then subjected to size-exclusion chromatography using a Superdex 200 10/300 Increase column pre-equilibrated in SE300 buffer (20 mM HEPES 7.4, 300 mM NaCl, 10 % glycerol, 1 mM TCEP). Pure fractions, as assessed by SDS-PAGE and Coomassie staining, were pooled and concentrated using a 30 kDa MWCO Amicon Ultra centrifugal filter. The protein concentration was calculated based on the absorbance at 280 nm and the predicted molar extinction coefficient (DNAstar Lasergene Suite). Aliquots were frozen in liquid nitrogen and stored at −80 °C.

6xHis-GST-TEV-yCTD was purified following a similar scheme as described earlier ^47^ with the following modifications. The clarified extract was applied to a 5 ml HisTrap HP column equilibrated in lysis buffer LB150. The column was extensively washed using high salt buffer HSB800 (20 mM HEPES pH 7.4, 800 mM NaCl, 30 mM imidazole, 10 % glycerol, 1 mM DTT, 0.284 μg/ml leupeptin, 1.37 μg/ml pepstatin A, 0.17 mg/ml PMSF, 0.33 mg/ml benzamidine) and equilibrated again in LB150. A pre-equilibrated 5 ml HiTrap Q HP column (GE healthcare) was attached in-line to the HisTrap column, which was subsequently eluted using a linear gradient from 0-100 % nickel elution buffer 150 (20 mM HEPES pH 7.4, 150 mM NaCl, 500 mM imidazole, 10 % glycerol, 1 mM DTT, 0.284 μg/ml leupeptin, 1.37 μg/ml pepstatin A, 0.17 mg/ml PMSF, 0.33 mg/ml benzamidine). The flow-through fractions were analyzed by SDS-PAGE and Coomassie staining, pooled and concentrated using a 30 kDa MWCO Amicon Ultra centrifugal filter unit. The sodium chloride concentration was adjusted to 50 mM and the protein was applied to a 1 ml HiTrap S column (GE healthcare). The flow-through was concentrated using a 30 kDa MWCO Amicon Ultra concentrator and then separated on an equilibrated Superdex 200 10/300 Increase column (GE healthcare) with buffer SE300. Individual fractions were analyzed by SDS-PAGE and Coomassie staining, pure fractions were pooled and concentrated with a 30 kDa MWCO Amicon Ultra filter unit. *E. coli* extract from the 6xHis-GST-TEV expression was applied to a 5 ml HisTrap HP column, washed with HSB800 and eluted with nickel elution buffer 150. The protein was concentrated using a 10 kDa MWCO Amicon filter unit and directly subjected to size-exclusion chromatography as described above. Concentrated protein solutions were aliquoted, flash-frozen in liquid nitrogen, and stored at −80 °C.

The recombinant *S. cerevisiae* TFIIH kinase module consisting of the three subunits Kin28, Ccl1 and Tfb3 was prepared as described ^48^. For the purification of the three-subunit human TFIIH kinase module (CDK7, cyclin H and Mat1), insect cells were lysed by sonication in lysis buffer (20 mM K-HEPES pH 7.0, 400 mM KCl, 10% glycerol, 1 mM MgCl_2_, 10 μM ZnCl_2_, 5 mM β-mercaptoethanol, 30 mM imidazole pH 8, 0.284 μg/ml leupeptin, 1.37 μg/ml pepstatin A, 0.17 mg/ml PMSF, 0.33 mg/ml benzamidine). Clarified cell lysate was applied onto a HisTrap HP 5 ml column (GE Healthcare), washed with 20 CV of lysis buffer and eluted with a linear gradient of 0-100% of elution buffer (20 mM K-HEPES pH 7, 400 mM KCl, 10% glycerol, 1 mM MgCl_2_, 10 μM ZnCl_2_, 5 mM β-mercaptoethanol, 500 mM imidazole pH 8, 0.284 μg/ml leupeptin, 1.37 μg/ml pepstatin A, 0.17 mg/ml PMSF, 0.33 mg/ml benzamidine) in 10 CV. Peak fractions were combined, supplemented with 2 mg of 6xHis-tagged TEV protease and dialyzed overnight against 2 L dialysis buffer (20 mM K-HEPES pH 7, 400 mM KCl, 10% glycerol, 1 mM MgCl_2_, 10 μM ZnCl_2_, 5 mM β-mercaptoethanol). The dialyzed solution was applied onto a HisTrap HP 5 ml column preequilibrated in dialysis buffer. The trimeric complex was eluted with 10% elution buffer and concentrated using an Amicon Ultra 15 ml 30 kDa MWCO centrifugal concentrator. The sample was applied to a Superdex 200 10/300 GL size exclusion column (GE healthcare) pre-equilibrated in storage buffer (20 mM K-HEPES pH 7, 350 mM KCl, 10% glycerol, 1 mM MgCl_2_, 10 μM ZnCl_2_, 5 mM β-mercaptoethanol). Peak fractions containing stoichiometric kinase trimer were pooled, concentrated using an Amicon Ultra 15 ml 30 kDa MWCO centrifugal concentrator to 130 μM, aliquoted, flash frozen in liquid nitrogen and stored at −80 °C.

The identity of all purified proteins was confirmed by LC-MS/MS analysis.

**Pol II preparation and fluorescent labeling.** Pol II was prepared from the *S. cerevisiae* strain BJ5464 as described ^49^ and treated with lambda phosphatase during purification. The Pol II subunit RPB3 contains an N-terminal biotin acceptor peptide, which can be biotinylated *in vitro* by the bacterial biotin-protein ligase BirA and used for site-specific labeling with fluorescent streptavidin conjugates. For this, 200 μg Pol II were incubated with 6 μg BirA, 100 μM D(+)-biotin and 2 mM ATP for 2 h at 20 °C in Pol II buffer (10 mM HEPES pH 7.2, 200 mM KCl, 5 % glycerol, 2 mM DTT). Excess biotin was removed using a Micro Bio-Spin 6 column (Biorad) according to the manufacturer’s suggestions. A small fraction of biotinylated Pol II was bound to streptavidin-coupled Dynabeads M-280 (Thermo Fisher Scientific) to confirm quantitative biotinylation. The remaining biotinylated Pol II was reacted with Alexa 594-coupled streptavidin (Thermo Fisher Scientific, ~2x molar excess) for 20 min at 20 °C. Pol II was then separated from unbound streptavidin by size-exclusion chromatography using a Superose 6 10/300 column (GE healthcare) equilibrated in Pol II buffer. Pol II-containing fractions were pooled, concentrated (100 kDa MWCO Amicon Ultra spin filter unit), and flash-frozen aliquots were stored in the dark at −80 °C.

**CTD phosphorylation.** GST-yCTD was phosphorylated using the recombinant *S. cerevisiae* TFIIH kinase module. For this, 50 μM GST-yCTD were incubated with 0.4 μM kinase module and 3 mM ATP for 1 h at 30 °C in kinase reaction buffer (20 mM HEPES pH 7.4, 200 mM NaCl, 5 mM MgCl_2_, 10 % glycerol, 1 mM TCEP). Upon completion, the phosphorylation reaction was quenched by addition of EDTA to a final concentration of 10 mM. Phosphorylation of MBP-hCTD was performed using the recombinant human TFIIH kinase module. For this, MBP-hCTD (100 μM) was incubated with 2 μM kinase module in reaction buffer (20 mM HEPES pH 7.4, 260 mM NaCl, 20 mM MgCl_2_, 20 μM ZnCl_2_, 10 % glycerol, 2 mM TCEP). The reaction was started by addition of 8 mM ATP, incubated for 1 h at 30 °C and quenched by addition of 40 mM EDTA. Control reactions lacking either the kinase or ATP were conducted in both cases under identical conditions. After completion of GST-yCTD and MBP-hCTD phosphorylation experiments, all reactions were mixed with 20 % dextran (in buffer containing 20 mM HEPES pH 7.4, 200 mM NaCl) at a ratio of 1:4 (vol/vol) and then analyzed microscopically (as described below). To study phosphorylation-induced dissolution of preformed CTD droplets, MBP-hCTD was mixed at a final concentration of 20 μM into 16 % dextran containing 20 mM HEPES pH 7.4, 220 mM NaCl, 1.6 mM ATP, 4 mM MgCl_2_, 20 μM ZnCl_2_, and 1 mM TCEP to induce phase-separation. Immediately before imaging, the reaction was started by addition of human TFIIH kinase module to a final concentration of 0.4 μM and immediately analyzed by microscopy.

**Kinase activity assay.** Kinase activity was analyzed by mobility shift assays. One microgram CTD fusion protein from kinase and control reactions was separated on 4-15% tris-glycine Protean TGX polyacrylamide gels (Biorad) and stained with Coomassie solution (InstantBlue, Expedeon). Phosphorylation of the CTD substrates by human and yeast TFIIH kinase modules results in a pronounced decrease of electrophoretic mobility. Phosphorylation of the CTD residue Ser5 was confirmed by immunoblotting. For this, samples (100 ng/lane) were separated on 4-15 % tris-glycine Protean TGX gels and blotted onto a PVDF membrane with a Trans-Blot Turbo Transfer System (Biorad). The membrane was blocked for 1-2 h at room temperature with 5 % (w/v) milk powder in phosphate-buffered saline containing 0.1 % Tween-20 (PBST). The blocked membrane was then incubated with either anti-MBP HRP conjugate (ab49923; Abcam) or anti-GST HRP conjugate (RPN1236; GE healthcare) for 2 h at room temperature. SuperSignal West Pico Chemiluminescent Substrate (Thermo Fisher) was used to develop the membrane before scanning with a ChemoCam Advanced Fluorescence imaging system (Intas Science Imaging). For immunoblot analysis of CTD phosphorylation, the membrane was subsequently stripped by incubation in stripping buffer (200 mM glycine-HCl pH 2.2, 0.1 % SDS, 1 % Tween-20), blocked with 5 % (w/v) milk powder in PBST, and probed overnight at 4 °C with primary CTD antibody against phosphorylated Ser5 (3E8; diluted 1:60 in 2.5 % (w/v) milk powder in PBST). The anti-Ser5 CTD antibody was a kind gift of Dirk Eick (Molecular Epigenetics Research Unit, Helmholtz Center, Munich). The membrane was then incubated with an anti-rat HRP-conjugate (A9037, Sigma-Aldrich) in 2.5 % milk-PBST for 1 h at room temperature and developed as describe above.

**Disorder prediction.** Recent cryo-EM analysis of mammalian RNA polymerase II could derive an atomic model only to RPB1 position P1487^50^, indicating a high conformational flexibility of the following RPB1-linker and the C-terminal repeat domain. We thus used the VLXT predictor implemented in PONDR^51^ to calculate the disorder propensity for the human RPB1 residues 1488-1970.

**CD spectroscopy.** Far-UV CD measurements were performed on a Chirascan spectrometer (Applied Photophysics, Ltd) at 25 °C using a 0.2 mm path length cuvette. The concentration of MBP-hCTD and MBP-yCTD was 5 μM in 20 mM NaPi, pH 7.4. CD spectra were recorded from 180 to 280 nm with an integration time of 0.5 s, and experiments were repeated three times. The spectra of human and yeast CTD were obtained through subtraction of the spectrum of MBP and correction of the baseline using buffer. Data are expressed in terms of the mean residual ellipticity (*θ*) in [deg/(cm^2^ dmol)].

**NMR spectroscopy.** Peptides comprising one (1R-CTD; 7 residues), two (2R-CTD; 14 residues) and three (3R-CTD; 21 residues) YSPTSPS-repeats were synthesized by GenScript with acetyl- and amide protection groups at the N- and C-termini, respectively. NMR spectra were recorded at 5 °C on Bruker 600 and 700 MHz spectrometers with triple-resonance cryogenic probes for 1.0 mM 3R-CTD peptide (20 mM HEPES, pH 7.4, 200 mM NaCl, and 90% H_2_O/10% D_2_O) and 0.5 mM phosphorylated 3R-CTD peptide. For phosphorylation, 3R-CTD was incubated with 2 μM TFIIH kinase at 37 °C during 18h (20 mM HEPES, pH 7.4, 200 mM NaCl, 3 mM MgCl_2_, 3 mM ATP, and 90% H_2_O/10% D_2_O). Spectra were processed with the software TopSpin (Bruker) and analyzed using CCPN Analysis ^52^. Sequence-specific backbone and side-chain resonance assignments of nonphosphorylated and phosphorylated 3R-CTD peptide were achieved through ^1^H-^15^N heteronuclear single quantum coherence (HSQC) and ^1^H-^13^C HSQC experiments at natural abundance, together with 2D ^1^H-^1^H TOCSY (100 ms mixing time) and 2D ^1^H-^1^H NOESY (200 and 300 ms mixing time) experiments. Resonance assignments of 3R-CTD were further validated through comparison with NMR spectra recorded for 1R-CTD and 2R-CTD peptides.

**Differential Interference Contrast (DIC) and fluorescence microscopy.** Droplet formation of protein samples was monitored by DIC and fluorescence microscopy. Samples were fluorescently labeled using Alexa Fluor 488 Microscale Protein Labeling Kit (Thermo Fisher Scientific, #A30006) according to manufacturer’s instructions. Small amounts (<0.5 μM) of labeled protein, which are not sufficient to induce droplet formation by itself, were mixed with unlabeled protein to the final concentration indicated in the text. In experiments with Ficoll PM 400 (Sigma, #F4375) at the final concentration of 150 mg/ml, buffer containing 20 mM HEPES, 200 mM NaCl, pH 7.4 was used. In experiments using dextran T500 (Pharmacosmos) as a crowding agent, dextran was added to reach the indicated final concentrations in 20 mM HEPES, 220 mM NaCl, pH 7.4. 5-10 μl of samples were loaded onto glass slides, covered with ø 18 mm coverslips and sealed. DIC and fluorescent images were acquired on a Leica DM6000B microscope with a 63x objective (water immersion) and processed using the FIJI software (NIH). In experiments requiring MBP-tag removal, fusion proteins were incubated with TEV protease in molar ratio TEV:protein=1:25 for 3 h at 25 °C. Complete tag removal was confirmed by SDS-PAGE analysis and Coomassie staining of the digested samples.

In experiments with aliphatic alcohols, the MBP-tag was cleaved off from MBP-yCTD and MBP-hCTD as indicated above, followed by addition of the protein to a premix containing dextran (final concentration 16%) and either 1,6-hexanediol (Sigma, #240117) or 2,5-hexanediol (Sigma, #H11904). The final protein concentration in the sample was 50 μM for yCTD and 20 μM for hCTD and hexanediol concentrations varied from 2.5 to 10%. Samples were imaged by DIC microscopy as indicated above.

All experiments with droplet formation were performed at room temperature except when the influence of temperature was tested. In the later case, MBP-hCTD or MBP-yCTD was mixed with small amounts (<0.2 μM) of the corresponding Alexa 488-labeled protein, from which the MBP-tag was cleaved off using TEV protease as described above. Final protein concentrations in the samples were 20 μM for MBP-hCTD and 40 μM for MBP-hCTD in 20 mM HEPES, 220 mM NaCl, pH 7.4 with 16% dextran. Samples were then incubated for one hour on ice (4 °C), room temperature (22 °C) or in an incubator at 37 °C or 45 °C before microscopy analysis. Labeled (without MBP-tag) and unlabeled (MBP-tagged) proteins were also mixed in experiments testing the influence of ionic strength. Final protein concentrations were 10 μM for MBP-hCTD and 40 μM for MBP-yCTD and samples contained indicated NaCl concentrations in 20 mM HEPES, pH 7.4 containing 16% dextran.

**Co-recruitment experiments.** For CTD co-recruitment experiments, droplets were made with 20 μM MBP-hCTD or GST-yCTD in 20 mM HEPES, 220 mM NaCl, pH 7.4 containing 16% dextran. Droplets were visualized through addition of 0.6 μM of tetramethylrhodamin (TMR)-labeled peptide with the sequence YSPTSPS, i.e. corresponding to one consensus heptad repeat. Subsequently, small amounts (< 0.5 μM) of Alexa 488 labeled GST-yCTD or MBP-hCTD were added to pre-formed MBP-hCTD or GST-yCTD droplets, respectively. Co-recruitment was assessed by imaging on a Leica DM6000B microscope as described above using DIC in combination with red and green channels for fluorescence (GFP and N3 filter cubes).

For Pol II co-recruitment experiments, Alexa 594-labeled Pol II (final concentration 0.02 μM) was mixed with pre-formed GST-yCTD droplets (final concentration of 25 μM) that were visualized by addition of Alexa 488-labeled GST-yCTD (final concentration of 2.3 μM) in 20 mM HEPES, 220 mM NaCl, pH 7.4 containing 16% dextran. Co-recruitment was documented by DIC and fluorescent microscopy using red and green channels (GFP and N3 filter cubes) on a Leica DM6000B microscope as described.

***In vitro* FRAP experiments.** The dynamics of human and yeast CTD molecules in the phase-separated state were investigated by fluorescence recovery after photobleaching (FRAP). MBP-tagged human and yeast CTD proteins were labeled on a single Cys residue that is present C-terminal to the TEV protease cleavage site (see above) using Alexa Fluor 488 C5 Maleimide dye (Thermo Fisher Scientific, #A10254) according to the manufacturer’s recommendations. Briefly, proteins were incubated in a light protected Eppendorf tube with the dye freshly dissolved in DMSO in a molar ratio 1:15=protein:dye in 20 mM HEPES pH 7.4, 300 mM NaCl, 1 mM TCEP, 10% glycerol for 3 h at room temperature. Excess label and salt were removed by desalting samples twice with 0.5 mL 7000 MWKO Zeba spin desalting columns (Thermo Fisher Scientific, #89882). The MBP-tag was then cleaved from both labeled and unlabeled human and yeast CTD using TEV protease as indicated above. Droplets for FRAP measurements were made in 16% dextran T500 in 20 mM HEPES, 220 mM NaCl, pH 7.4 by adding mixtures of labeled and unlabeled yCTD (or hCTD) in a molar ratio of 1:100 to the final CTD concentration of 20 μM. To minimize droplet movement, FRAP recordings were done after approx. 30 minutes, which is the time required for freshly formed droplets to settle down on the glass slide and become less mobile.

FRAP experiments were recorded on a Leica TCS SP8 confocal microscope using 63x objective (water immersion) at a zoom corresponding to a pixel size of 96 nm × 96 nm and using the 488 argon laser line. A circular region of approx. 1.4 μm in diameter was chosen in a region of homogenous fluorescence away from the droplet boundary and bleached with 10 iterations of full laser power. Recovery was imaged at low laser intensity (0.057%). 50 frames were recorded with one frame per 330 ms. Pictures were analyzed in FIJI (NIH) and FRAP recovery curves were calculated using standard methods. For calculating half time recoveries, normalized values from each recording were separately fit to a single exponential model and half time recoveries were presented as mean ± standard error.

**Cell line establishment and characterization.** Human U2OS osteosarcoma cells (Research Resource Identifier: RRID:CVCL_0042) were grown in a Sanyo copper alloy IncuSafe humidified incubator (MCO-18AIC(UV)) at 37°C/5.5% CO2 in low glucose DMEM with 10% FBS (full recipe: 500 mL DMEM (ThermoFisher #10567014), 50 mL fetal bovine serum (HyClone FBS SH30910.03 lot #AXJ47554) and 5 mL Penicillin-streptomycin (ThermoFisher #15140122)) and were passaged every 2-4 days before reaching confluency. Plasmids expressing N-terminally tagged (either Dendra2 or Halo) α-amanitin resistant mutated (N792D) human RPB1 were stably transfected into U2OS cells using Fugene 6 following the manufacturer’s instruction (Promega #E2692). The RPB1-52R vectors encode the 52 CTD repeats originally present in the endogenous RPB1 cDNA. The RPB1-25R expressing vectors contain only 25 repeats out of the 52 repeats corresponding to repeats 1 to 21 and repeats 49 to 52. The RPB1-70R cell lines are expressing either a Dendra2-RPB1 protein containing 66 repeats in its CTD (repeats 1 to 51, then repeats 38 to 52) or an Halo-RPB1 protein containing 70 repeats in its CTD (repeats 1 to 47, then repeats 42 to 47 then repeats 38 to 52) as assessed by sequencing of the *RPB1* mRNA expressed in these cells. Details of cloning strategies are available upon request. α-amanitin (SIGMA #A2263) was used during the stable selection process at a concentration of 2 μg/mL and was used thereafter in permanence in the culture of the cells at a concentration of 1 μg/mL to avoid endogenous RPB1 reexpression as described in ^3^. Even though these lines cannot genotypically be considered as endogenously tagged (the endogenous wild-type *RPB1* gene is still present, a cDNA expressing the tagged version of RPB1 is incorporated in the genome), phenotypically they can as the expression of endogenous RPB1 protein is replaced by the tagged version of the protein at all time.

RT-PCR analysis (Superscript III with oligo (dT)_20_, Invitrogen (#18080093) and NEB Phusion^®^ High-Fidelity DNA Polymerase (#M0530S) followed by sequencing was performed to confirm the sequence of the RPB1-CTD expressed in the various cell lines (more details of all the molecular biology characterizations available upon request).

**Western blot.** Cells were collected after ice-cold PBS wash by scraping into 0.5 mL/10 cm plate of high-salt lysis buffer (0.5 M NaCl, 50 mM HEPES, 5 mM EDTA, 0.5% NP-40 and protease inhibitors), with 125 U/mL of benzonase (Novagen, EMD Millipore), passed through a 25G needle, rocked at 4°C for 30 min, centrifuged at maximum speed at 4°C for 20 min. Supernatants were quantified by Bradford method. Same amount of proteins were loaded onto 7% Bis-Tris SDS-PAGE gel, transferred onto nitrocellulose membrane (Amershan Protran 0.45 um NC, GE Healthcare) for 2 hr at 80V, blocked in TBS-Tween with 5% milk for at least 1 h at room temperature and blotted overnight at 4°C with primary antibodies (anti-Pol II (N20) from SantaCruz #sc-899, anti-LaminA from Abcam #ab26300) in TBS-T with 5% milk. HRP-conjugated secondary antibodies were diluted 1:5000 in TBS-T with 5% milk and incubated at room temperature for one hour.

**FACS analysis.** Expression of the exogenous RPB1 protein was assessed by flow cytometry analysis on live cells on a BD LSRFortessa, performed according to the manufacturer’s protocols. For the Halo-tagged line, Halo-TMR labeling (500 nM) was performed for 30mn at 37°C before harvesting the cells.

**xCELLigence analysis.** The Cell Index (a representation of cell growth and viability) was measured in real time using the RTCA-SP (Acea Biosciences) according to manufacturer’s instructions. Cells were seeded at a density of 4000 cells/well. The Cell Index was normalized at 3h after seeding to account for slight variations in the number of counted cells between various lines.

**Doubling Time Analysis.** Doubling time analysis was performed (using FarRed CFSE from CellTrace™ CFSE Cell Proliferation Kit, ThermoFisher Scientific #C34554) to compare the growth capacity of the different lines. More precisely for doubling time analysis, data was collected on a BD Bioscience LSR Fortessa; geometric fluorescent mean intensity of each sample for each time point (day 1 to day 5) was extracted from FlowJo and the average change over the 5 day period was calculated. The average change was then converted to log scale to calculate the doubling time per 24 hours.

**Cell imaging conditions.** For live-cell imaging, the medium was identical except DMEM without phenol red was used (ThermoFisher #31053028). U2OS cells expressing α-amanitin-resistant Halo-RPB1-25R, Halo-RPB1-52R or Halo-RPB1-70R were grown overnight with α-amanitin on plasma-cleaned 25 mm circular no 1.5H cover glasses (Marienfeld High-Precision 0117650). For the flavopiridol experiments cells were treated for 30-45 min with flavopiridol (2 μM final concentration) then imaged for a maximum of 30-45 min. Prior to all experiments, the cover glasses were plasma-cleaned and then stored in isopropanol until use. For live-cell FRAP experiment, cell preparation was identical except cells where grown on glass-bottom (thickness #1.5) 35 mm dishes (MatTek P35G-1.5-14-C).

**PALM imaging.** Six movies of ~ 50,000 frames were acquired for each condition at 30 ms/frame. The axial drift was corrected in real time with a perfect focus system. A cylindrical lens was added to the system to induce astigmatism in the point-spread function (PSF) of the optical setup. 300,000 detections were collected on average per movie. Single molecule detection and localization was performed with a modified version of the multiple-target tracking algorithm. The 3D position of single detections was inferred from the lateral elongation of the PSF. The lateral drift of the sample was corrected by using fluorescent beads (TetraSpeck microspheres). To correct for blinking of the Dendra2 fluorophore, detections in a disk of 30 nm and adjacent in time were grouped and averaged.

Nuclei and nucleoli were automatically detected and segmented for further processing. N(r) is the estimate of the expected number of neighbors within a distance r of a given point of the sample:

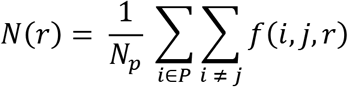

where P is the set of all detections and N_p_ the total number of detections. The *f* function ^53 54^ corrects, for biases generated by points located at short distances to the borders (nucleus or nucleoli):

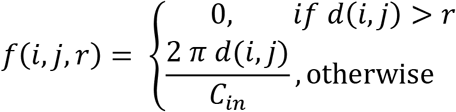

where *d(i,j)* is the distance between *i* and j and *C*_*in*_ the length of the part of the circle of radius *d(i,j)* centered on *i* which is inside the area of study, the nucleus.

The null hypothesis, complete spatial randomness (CSR) is a homogeneous Poisson process with intensity λ, equal to the density of detection in the area of study A: 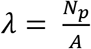

We estimated four spatial statistics based on N(r): n(r), K(r), L(r) and G(r) ^54,55^. The local neighbor density function, is defined as *n*(*r*) = *N*(*r*) / *πr*^2^. The K-Ripley function is defined as *K*(*r*) = *N*(*r*) / *λ*. The linearized K-Ripley function is given by 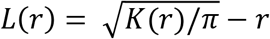. The pair density function G(r) is simply the derivative of the K(r).

Under CSR, the expected value taken by n(r) (resp. K(r), L(r) and G(r)) is λ (resp. *πr*^2^, 0 and 1). The triangulation of the areas was performed with a custom python script and we used the ADS R package ^56^ to estimate the four spatial statistics. In order to estimate the standard deviation and standard error associated with these measurements we performed a bootstrapping analysis of the data set. We randomly selected 10,000 detections from each original data set a 100 times and feed these sub sampled data set to the R script computing the spatial statistics.

**Single-molecule imaging (spaSPT).** After overnight growth, cells were labeled with 50 nM PA-JF_549_ ^57^ for ~15-30 min and washed twice (one wash: medium removed; PBS wash; replenished with fresh medium). At the end of the final wash, the medium was changed to phenol red-free medium keeping all other aspects of the medium the same (and adding back α-amanitin). Singlemolecule imaging was performed on a custom-built Nikon TI microscope equipped with a 100x/NA 1.49 oil-immersion TIRF objective (Nikon apochromat CFI Apo TIRF 100x Oil), EM-CCD camera (Andor iXon Ultra 897; frame-transfer mode; vertical shift speed: 0.9 μs; −70°C), a perfect focusing system to correct for axial drift and motorized laser illumination (Ti-TIRF, Nikon), which allows an incident angle adjustment to achieve highly inclined and laminated optical sheet illumination ^58^. An incubation chamber maintained a humidified 37°C atmosphere with 5% CO_2_ and the objective was also heated to 37°C. Excitation was achieved using a 561 nm (1 W, Genesis Coherent) laser for PA-JF_549_. The excitation laser was modulated by an acousto-optic tunable filter (AA Opto-Electronic, AOTFnC-VIS-TN) and triggered with the camera TTL exposure output signal. The laser light was coupled into the microscope by an optical fiber and then reflected using a multi-band dichroic (405 nm/488 nm/561 nm/633 nm quad-band, Semrock) and then focused in the back focal plane of the objective. Fluorescence emission light was filtered using a single bandpass filter placed in front of the camera using the following filters: Semrock 593/40 nm bandpass filter. The microscope, cameras, and hardware were controlled through NIS-Elements software (Nikon).

We recorded single-molecule tracking movies using our previously developed technique, stroboscopic photo-activation Single-Particle Tracking (spaSPT) ^34,36^. Briefly, 1 ms 561 nm excitation (100% AOTF) of PA-JF_549_ was delivered at the beginning of the frame to minimize motion-blurring; 405 nm photo-activation pulses were delivered during the camera integration time (~447 μs) to minimize background and their intensity optimized to achieve a mean density of ~1 molecule per frame per nucleus. 30,000 frames were recorded per cell per experiment. The camera exposure time was 7 ms resulting in a frame rate of approximately 134 Hz (7 ms + ~447 μs per frame).

spaSPT data was analyzed (localization and tracking) and converted into trajectories using a custom-written Matlab implementation of the MTT-algorithm ^59^ and the following settings: Localization error: 10^−6.25^; deflation loops: 0; Blinking (frames): 1; max competitors: 3; max *D* (μm^2^/s): 20.

We recorded ~5-10 cells per replicate and performed three independent replicates on three different days. Specifically, across three replicates we imaged 29 cells for 25R Halo-RPB1 and obtained 448,362 trajectories with 690,682 unique displacements at a mean density of 1.2 localizations per frame. Similarly, we imaged 30 cells for 52R Halo-RPB1 and obtained 324,928 trajectories with 619,247 unique displacements at a mean density of 1.1 localizations per frame. Finally, we imaged 26 cells for 70R Halo-RPB1 and obtained 333,720 trajectories with 571,345 unique displacements at a mean density of 1.0 localizations per frame. In the flavopiridol treated experiment, we imaged 13 cells for 25R Halo-RPB1 and obtained 598,941 trajectories with 926,057 unique displacements at a mean density of 2.4 localizations per frame. We imaged 15 cells for 52R Halo-RPB1 and obtained 395,206 trajectories with 671,492 unique displacements at a mean density of 1.5 localizations per frame. Finally, we imaged 28 cells for 70R Halo-RPB1 and obtained 616,088 trajectories with 1,030,523 unique displacements at a mean density of 1.9 localizations per frame.

**Model-based analysis of single-molecule tracking data using Spot-On.** To analyze the spaSPT data, we used our previously described kinetic modeling approach (Spot-On) ^34,36^. Briefly, we analyze each replicate separately and the reported bound fractions and free diffusion coefficients are reported as the mean +/− standard deviation from analyzing each replicate separately. We merged the data from all cells (~9-10) for each replicate, compiled histograms of displacements and then fit the displacement cumulative distribution functions for 7 time points using a two-state model that assumes that Pol II can either exist in an immobile (e.g. chromatin-associated) or freely diffusive state:

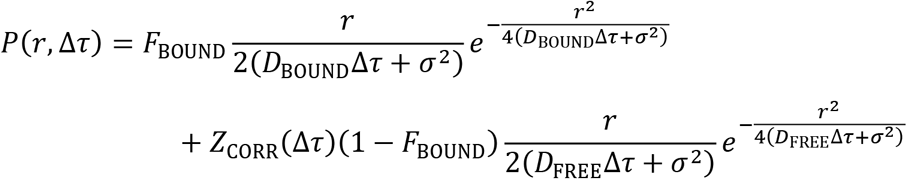

where:

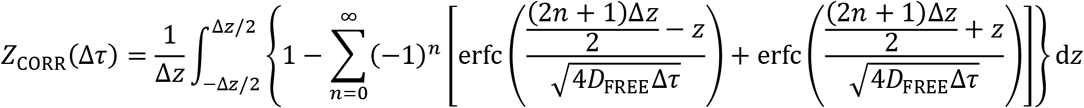

and:

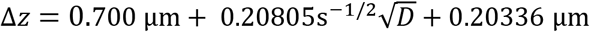

Here, *F*_BOUND_ is the fraction of molecules that are bound to chromatin, *D*_BOUND_ is the diffusion coefficient of chromatin bound molecules, *D*_FREE_ is the diffusion coefficient of freely diffusing molecules, *r* is the displacement length, Δ*τ* is lag time between frames, Δ*z* is axial detection range, *σ* is localization error (35 nm) and *Z*_CORR_ corrects for defocalization bias (i.e. the fact that freely diffusion molecules gradually move out-of-focus, but chromatin bound molecules do not). Model fitting and parameter optimization was performed using a non-linear least squares algorithm (Levenberg-Marquardt) implemented in the Matlab version of Spot-On (v1.0; GitLab tag 92cdf210) and the following parameters: dZ=0.7 μm; GapsAllowed=1; TimePoints: 7; JumpsToConsider=4; ModelFit=2; NumberOfStates=2; FitLocError=0; D_Free_2State=[0.4;25]; D_Bound_2State=[0.00001;0.05];

**Diffusion coefficient calculations.** The observed free diffusion coefficients obtained from fitting the spaSPT data with the Spot-On model (Brownian motion) were 3.74 +/− 0.178 μm^2^/s, 2.97 +/− 0.0912 μm^2^/s and 2.34 +/− 0.049 μm^2^/s for the 25R, 52R and 70R versions of Halo-Rpb1, respectively (mean +/− standard error). Given that the molecular weight of e.g. 25R is lower, one would expect the diffusion coefficient to be higher. To estimate whether this large difference could be explained by size alone or whether it might be due to reduced multivalent interactions, we consider the Stokes-Einstein relation according to which the diffusion coefficient is given by:

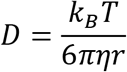

where *k*_*B*_ is Boltzmann’s constant, *T* is the absolute temperature, *η* is the viscosity of the liquid (the nucleoplasm here; assumed to be the same for both 25R, 52R and 70R) and *r* is the radius. The Stokes-Einstein equation assumes the particle to be a sphere and accordingly the radius is given by the volume, V:

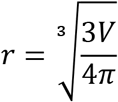

In turn, the volume is related to the mass, *m*, and density, *ρ*:

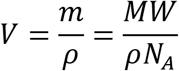

where *N*_*A*_ is Avogadro’s constant and *MW* is the molecular weight in atomic mass units (Daltons). Thus, the diffusion coefficient is related to the molecular weight by:

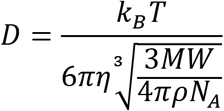

Thus using 25R and 52R as the example, the ratio between the diffusion coefficients of 25R and 52R Halo-Rpb1 (assuming that the density is the same) is:

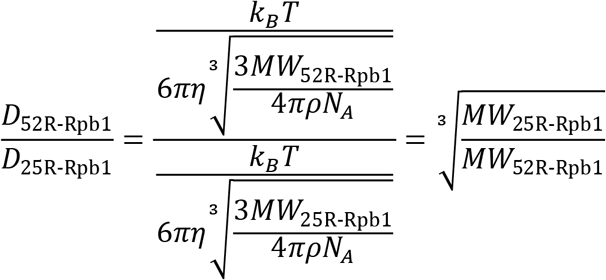

According to UniProt (P24928) the molecular weight of wild-type Rpb1 is 217.2 kDa (52R). The molecular weight of the HaloTag is 33.6 kDa. Thus, the molecular weight of Halo-Rpb1-52R is ~250.8 kDa, the molecular weight of Halo-Rpb1-25R is ~230.9 kD and the molecular weight of Halo-Rpb1-70R is ~258.1 kD. Thus, the expected difference in diffusion coefficients is:

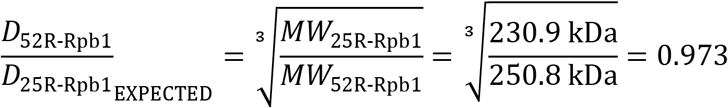

If we compare this to the experimentally observed ratio:

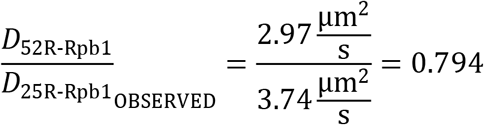

It becomes clear that size/mass difference alone cannot explain the large difference in diffusion coefficient that we observe in cells. To be comprehensive, below we list the Stokes-Einstein expected and the observed diffusion coefficient ratios for all the combinations:

| Comparison | Stokes-Einstein expectation | Observed ratio |
| --- | --- | --- |
| 25R vs. 52R | 0.973 | 0.794 |
| 25R vs. 70R | 0.964 | 0.626 |
| 52R vs. 70R | 0.991 | 0.789 |

For all three combinations, the observed ratio cannot be explained by the change in size/mass. Instead this indicates a higher propensity of the full-length CTD to engage in intermolecular interactions. Moreover, in the above calculations we have just considered the change in the mass of Rpb1. In reality, Rpb1 is likely diffusing as part of the Pol II holocomplex, thus the relative difference due to the smaller CTD (e.g. ~20 kDa between 25R and 52R) is actually much smaller than the calculations using only Rpb1 would suggest and thus the expected difference in diffusion coefficients due to mass/size would be even much closer to 1. We conclude that the mass/size difference between the 25R, 52R and 70R Pol II enzymes cannot explain their observed differences in diffusion coefficients.

**FRAP in cells.** FRAP experiments were performed and analyzed as previously described ^36^. Briefly, FRAP was performed on an inverted Zeiss LSM 710 AxioObserver confocal microscope equipped with a motorized stage, a full incubation chamber maintaining 37°C/5% CO_2_, a heated stage, an X-Cite 120 illumination source as well as several laser lines. Halo-TMR was excited using a 561 nm laser. Images were acquired on a 40x Plan NeoFluar NA1.3 oil-immersion objective at a zoom corresponding to a 100 nm × 100 nm pixel size and the microscope controlled using the Zeiss Zen software. In FRAP experiments, 300 frames were acquired at either 1 frame per second allowing 20 frames to be acquired before the bleach pulse to accurately estimate baseline fluorescence. A circular bleach spot (*r* = 10 pixels) was chosen in a region of homogenous fluorescence at a position at least 1 μm from nuclear and nucleolar boundaries. The spot was bleached using maximal 561 nm laser intensity and pixel dwell time corresponding to a total bleach time of ~1 s. We generally collected data from 3-5 cells per cell line per condition per day and all presented data is from at least 3 independent replicates on different days.

To quantify and drift-correct the FRAP movies, we used a previously described custom-written pipeline in MATLAB ^36^. Briefly, we manually identify the bleach spot. The nucleus is automatically identified by thresholding images after Gaussian smoothing and hole-filling (to avoid the bleach spot as being identified as not belonging to the nucleus). We use an exponentially decaying (from 100% to ~85% (measured) of initial over one movie) threshold to account for whole-nucleus photobleaching during the time-lapse acquisition. Next, we quantify the bleach spot signal as the mean intensity of a slightly smaller circle (*r* = 0.6 μm), which is more robust to lateral drift. The FRAP signal is corrected for photobleaching using the measured reduction in total nuclear fluorescence (~15% over 300 frames at the low laser intensity used after bleaching) and internally normalized to its mean value during the 20 frames before bleaching. We correct for drift by manually updating a drift vector quantifying cell movement during the experiment. Finally, drift- and photobleaching corrected FRAP curves from each single cell were averaged to generate a mean FRAP recovery. We used the mean FRAP recovery in all figures and error bars show the standard error of the mean.

## DATA AVAILABILITY STATEMENT

The in vitro datasets generated during and/or analyzed during the current study are available from the corresponding author on reasonable request. The PALM and SPT data are available at http://doi.org/10.5281/zenodo.1188488. The raw spaSPT data is available in Spot-On readable CSV format in the form of single-molecule trajectories. The Spot-On Matlab code is available together with a step-by-step guide at Gitlab: https://gitlab.com/tjian-darzacq-lab/spot-on-matlab. For additional documentation, please see also the Spot-On website https://SpotOn.berkeley.edu and previous publications ^34,36^.

## REFERENCES

1 Banani, S. F., Lee, H. O., Hyman, A. A. & Rosen, M. K. Biomolecular condensates: organizers of cellular biochemistry. Nat Rev Mol Cell Biol 18, 285–298 (2017).

2 Boeynaems, S. et al. Protein Phase Separation: A New Phase in Cell Biology. Trends Cell Biol (2018).

3 Cisse, II et al. Real-time dynamics of RNA polymerase II clustering in live human cells. Science 341, 664–667 (2013).

4 Cook, P. R. The organization of replication and transcription. Science 284, 1790–1795 (1999).

5 Brangwynne, C. P. et al. Germline P granules are liquid droplets that localize by controlled dissolution/condensation. Science 324, 1729–1732 (2009).

6 Molliex, A. et al. Phase separation by low complexity domains promotes stress granule assembly and drives pathological fibrillization. Cell 163, 123–133 (2015).

7 Han, T. W. et al. Cell-free formation of RNA granules: bound RNAs identify features and components of cellular assemblies. Cell 149, 768–779 (2012).

8 Hnisz, D., Shrinivas, K., Young, R. A., Chakraborty, A. K. & Sharp, P. A. A Phase Separation Model for Transcriptional Control. Cell 169, 13–23 (2017).

9 Li, P. et al. Phase transitions in the assembly of multivalent signalling proteins. Nature 483, 336–340 (2012).

10 Martin, E. W. & Mittag, T. Relationship of Sequence and Phase Separation in Protein Low-Complexity Regions. Biochemistry (2018).

11 Csizmok, V., Follis, A. V., Kriwacki, R. W. & Forman-Kay, J. D. Dynamic Protein Interaction Networks and New Structural Paradigms in Signaling. Chem Rev 116, 6424–6462 (2016).

12 Zaborowska, J., Egloff, S. & Murphy, S. The pol II CTD: new twists in the tail. Nat Struct Mol Biol 23, 771–777 (2016).

13 Hsin, J. P. & Manley, J. L. The RNA polymerase II CTD coordinates transcription and RNA processing. Genes Dev 26, 2119–2137 (2012).

14 Meinhart, A., Kamenski, T., Hoeppner, S., Baumli, S. & Cramer, P. A structural perspective of CTD function. Genes Dev 19, 1401–1415 (2005).

15 Simonti, C. N. et al. Evolution of lysine acetylation in the RNA polymerase II C-terminal domain. BMC Evol Biol 15, 35 (2015).

16 West, M. L. & Corden, J. L. Construction and analysis of yeast RNA polymerase II CTD deletion and substitution mutations. Genetics 140, 1223–1233 (1995).

17 Gibbs, E. B. et al. Phosphorylation induces sequence-specific conformational switches in the RNA polymerase II C-terminal domain. Nat Commun 8, 15233 (2017).

18 Portz, B. et al. Structural heterogeneity in the intrinsically disordered RNA polymerase II C-terminal domain. Nat Commun 8, 15231 (2017).

19 Janke, A. M. et al. Lysines in the RNA Polymerase II C-Terminal Domain Contribute to TAF15 Fibril Recruitment. Biochemistry (2017).

20 Cagas, P. M. & Corden, J. L. Structural studies of a synthetic peptide derived from the carboxyl-terminal domain of RNA polymerase II. Proteins 21, 149–160 (1995).

21 Hyman, A. A., Weber, C. A. & Julicher, F. Liquid-liquid phase separation in biology. Annu Rev Cell Dev Biol 30, 39–58 (2014).

22 Burke, K. A., Janke, A. M., Rhine, C. L. & Fawzi, N. L. Residue-by-Residue View of In Vitro FUS Granules that Bind the C-Terminal Domain of RNA Polymerase II. Mol Cell 60, 231–241 (2015).

23 Kwon, I. et al. Phosphorylation-regulated binding of RNA polymerase II to fibrous polymers of low-complexity domains. Cell 155, 1049–1060 (2013).

24 Brangwynne, C. P., Tompa, P. & Pappu, R. V. Polymer physics of intracellular phase transitions. Nat Phys 11, 899–904 (2015).

25 Pak, C. W. et al. Sequence Determinants of Intracellular Phase Separation by Complex Coacervation of a Disordered Protein. Mol Cell 63, 72–85 (2016).

26 Kato, M. & McKnight, S. L. A Solid-State Conceptualization of Information Transfer from Gene to Message to Protein. Annu Rev Biochem (2017).

27 Kapust, R. B. & Waugh, D. S. Escherichia coli maltose-binding protein is uncommonly effective at promoting the solubility of polypeptides to which it is fused. Protein Sci 8, 1668–1674 (1999).

28 Darzacq, X. et al. In vivo dynamics of RNA polymerase II transcription. Nat Struct Mol Biol 14, 796–806 (2007).

29 Becker, M. et al. Dynamic behavior of transcription factors on a natural promoter in living cells. EMBO Rep 3, 1188–1194 (2002).

30 Cho, W. K. et al. Super-resolution imaging of fluorescently labeled, endogenous RNA Polymerase II in living cells with CRISPR/Cas9-mediated gene editing. Sci Rep 6, 35949 (2016).

31 Huang, B., Wang, W., Bates, M. & Zhuang, X. Three-dimensional super-resolution imaging by stochastic optical reconstruction microscopy. Science 319, 810–813 (2008).

32 Nicovich, P. R., Owen, D. M. & Gaus, K. Turning single-molecule localization microscopy into a quantitative bioanalytical tool. Nat Protoc 12, 453–460 (2017).

33 Annibale, P., Vanni, S., Scarselli, M., Rothlisberger, U. & Radenovic, A. Identification of clustering artifacts in photoactivated localization microscopy. Nat Methods 8, 527–528 (2011).

34 Hansen, A. S. et al. Robust model-based analysis of single-particle tracking experiments with Spot-On. Elife 7 (2018).

35 Sprague, B. L., Pego, R. L., Stavreva, D. A. & McNally, J. G. Analysis of binding reactions by fluorescence recovery after photobleaching. Biophys J 86, 3473–3495 (2004).

36 Hansen, A. S., Pustova, I., Cattoglio, C., Tjian, R. & Darzacq, X. CTCF and cohesin regulate chromatin loop stability with distinct dynamics. Elife 6 (2017).

37 Feaver, W. J., Svejstrup, J. Q., Henry, N. L. & Kornberg, R. D. Relationship of CDK-activating kinase and RNA polymerase II CTD kinase TFIIH/TFIIK. Cell 79, 1103–1109 (1994).

38 Morgan, D. O. Principles of CDK regulation. Nature 374, 131–134 (1995).

39 Conaway, J. W., Shilatifard, A., Dvir, A. & Conaway, R. C. Control of elongation by RNA polymerase II. Trends Biochem Sci 25, 375–380 (2000).

40 Scafe, C. et al. RNA polymerase II C-terminal repeat influences response to transcriptional enhancer signals. Nature 347, 491–494 (1990).

41 Chong, S. et al. Dynamic and Selective Low-Complexity Domain Interactions Revealed by Live-Cell Single Molecule Imaging. BioRxiv (2017).

42 Lu, H. et al. Phase-separation mechanism for C-terminal hyperphosphorylation of RNA polymerase II. Nature (2018).

43 Rice, P., Longden, I. & Bleasby, A. EMBOSS: the European Molecular Biology Open Software Suite. Trends Genet 16, 276–277 (2000).

44 Patel, A. et al. ATP as a biological hydrotrope. Science 356, 753–756 (2017).

45 Gradia, S. D. et al. MacroBac: New Technologies for Robust and Efficient Large-Scale Production of Recombinant Multiprotein Complexes. Methods Enzymol 592, 1–26 (2017).

46 Farnung, L., Vos, S. M., Wigge, C. & Cramer, P. Nucleosome-Chd1 structure and implications for chromatin remodelling. Nature 550, 539–542 (2017).

47 Battaglia, S. et al. RNA-dependent chromatin association of transcription elongation factors and Pol II CTD kinases. Elife 6 (2017).

48 Schilbach, S. et al. Structures of transcription pre-initiation complex with TFIIH and Mediator. Nature 551, 204–209 (2017).

49 Sydow, J. F. et al. Structural basis of transcription: mismatch-specific fidelity mechanisms and paused RNA polymerase II with frayed RNA. Mol Cell 34, 710–721 (2009).

50 Bernecky, C., Herzog, F., Baumeister, W., Plitzko, J. M. & Cramer, P. Structure of transcribing mammalian RNA polymerase II. Nature 529, 551–554 (2016).

51 Obradovic, Z. et al. Predicting intrinsic disorder from amino acid sequence. Proteins 53 Suppl 6, 566–572 (2003).

52 Vranken, W. F. et al. The CCPN data model for NMR spectroscopy: development of a software pipeline. Proteins 59, 687–696 (2005).

53 Goreaud, F. & Pelissier, R. On explicit formulas of edge effect correction for Ripley’s K-function. J Veg Sci 10, 433–438 (1999).

54 Ripley, B. D. Modeling Spatial Patterns. JR Stat Soc Series B Stat Methodol 39, 172–212 (1977).

55 Dixon, P. M. Ripley’s K function. (2002).

56 Pelissier, R. & Goreaud, F. Ads package for R: a fast unbiased implementation of the K-function family for studying spatial point patterns in irregular-shaped sampling windows. J Stat Softw 63, 1–18 (1999).

57 Grimm, J. B. et al. Bright photoactivatable fluorophores for single-molecule imaging. Nat Methods 13, 985–988 (2016).

58 Tokunaga, M., Imamoto, N. & Sakata-Sogawa, K. Highly inclined thin illumination enables clear single-molecule imaging in cells. Nat Methods 5, 159–161 (2008).

59 Serge, A., Bertaux, N., Rigneault, H. & Marguet, D. Dynamic multiple-target tracing to probe spatiotemporal cartography of cell membranes. Nat Methods 5, 687–694 (2008).

